# Making a clean break: contrasting leaf abscission dynamics across temperate leaf habits

**DOI:** 10.1101/2025.07.10.664177

**Authors:** Cade N Kane, Ian M Rimer, Scott AM McAdam

## Abstract

**Background and Aims:** Leaf senescence allows plants to reallocate nutrients from aging leaves, often storing them for future use. As chlorophyll breaks down, other pigments emerge, creating the vibrant colors of autumn. In many deciduous trees, senescence is followed by abscission the physical detachment of leaves at the abscission zone (AZ). Though closely linked, senescence and abscission are distinct; some plants senesce without abscising, while others abscise partially senesced leaves. Most research on these processes has focused on herbaceous and crop species, which do not naturally shed leaves.

**Methods:** To better understand how leaf senescence and abscission occur in temperate deciduous trees, we developed a novel method to quantify AZ competency (AZC) we also measured gas exchange, chlorophyll content, water potential, and abscisic acid in four tree species representing deciduous, brevi-deciduous, and marcescent leaf habits.

**Key results:** The two deciduous species showed contrasting patterns: one degraded chlorophyll and ceased photosynthesis before AZC developed; the other retained chlorophyll and continued photosynthesis until nearly all AZs became competent. The brevi-deciduous species lost most chlorophyll but developed AZs gradually over a longer period. The marcescent species fully senesced but did not develop AZC.

**Conclusions:** These findings show that senescence and abscission are distinct and variably timed processes across temperate tree species.

## Introduction

The temperate deciduous phenology of trees significantly influences the annual global carbon cycle, local ecosystems, and human cultural practices (Archetti et al., 2013; Knorr, 2000; Lim et al., 2007; Mittermeier et al., 2019; Richardson et al., 2010; Sandifer et al., 2015). These trees are typically most productive during the spring and summer, with productivity declining in autumn in preparation for winter (Bassow and Bazzaz, 1998). This preparation, termed senescence, involves the reabsorption of leaf nitrogen and magnesium as well as lipids and proteins, for storage in woody tissues (Maillard et al., 2015), including the degradation of chlorophyll, which reveals the vibrant foliar colors associated with autumn. In herbaceous species senescence is considered the final stage of leaf development (Buchanan-Wollaston, 1997; Lim et al., 2007; Smart, 1994). In deciduous trees, senescence is typically followed by active leaf abscission, the process by which the senescent leaf is fully detached from the branch which persists in dormancy (Li and Su, 2024). Most modern work on organ abscission focuses on reproductive organs in annuals (Li and Su, 2024). The study of the mechanism of autumnal senescence and abscission in trees remains underexplored (Gallinat et al., 2015). Yet, the physiological mechanisms regulating senescence and abscission in deciduous trees are critical for predicting how these processes may shift forest responses to climate change (Norby, 2021; Zani et al., 2021; Zohner et al., 2023).

The evolutionary origins of deciduous leaf habits are debated, but it is likely deciduousness evolved independently dozens of times (Edwards et al., 2017; Tiffney and Manchester, 2001) during the Late Cretaceous in response to drought in the tropics (Axelrod, 1966), and during the Late Miocene in response to radiation into Northern Latitudes (Pound et al., 2011; Utescher and Mosbrugger, 2007; Wolfe, 1987; Wolfe and Upchurch, 1986). This diversity in evolutionary origin suggests that there might be considerable mechanistic diversity in the regulators of senescence and abscission across seed plants. In temperate tree species, the primary environmental cues for autumnal leaf senescence are day length and temperature (Lim et al., 2007; Noodén et al., 1997; Noodén and Leopold, 1988). Species differ in sensitivity to these cues, some respond more strongly to decreasing temperatures, while others are more sensitive to shorter day lengths (Lang et al., 2019; Mariën et al., 2022; Michelson et al., 2018).

A suite of hormones are known to contribute to senescence, with ethylene widely regarded as a primary internal trigger and accelerator of leaf senescence (Aharoni and Lieberman, 1979; Iqbal et al., 2017; Jibran et al., 2013). Ethylene triggers a cascade of senescence associated genes (SAGs) (Breeze et al., 2011; Gan and Amasino, 1997), leading to the breakdown and remobilization of cellular components for storage or reuse elsewhere in the plant (Koyama, 2014). Another hormone, abscisic acid (ABA), is also considered a significant regulator of leaf senescence (Noodén and Leopold, 1988). Most of the evidence supporting a link between ABA and leaf senescence comes from studies on leaf discs, detached leaves, or experiments involving exogenous ABA application (Chin and Beevers, 1970; Gao et al., 2016; Gepstein and Thimann, 1980; Zakari et al., 2020). The role of ABA in natural leaf senescence remains unclear, while some studies report increasing ABA levels as leaves age (El-Antably et al., 1967; He and Jin, 1999), others have found that ABA levels actually decrease during senescence (Uzelac et al., 2016), with no difference in senescence rates between wild-type and ABA-deficient sunflowers (McAdam et al., 2022). In trees, increased ABA levels resulting from stem girdling which disrupts phloem flow can hasten leaf senescence (Kane and McAdam, 2023; Lihavainen et al., 2021; López et al., 2015; Mitchell et al., 2017; Setter et al., 1980). Similarly, the application of exogenous ABA has been shown to accelerate senescence (Dann et al., 1984; Guak and Fuchigami, 2001; Lihavainen et al., 2021). Despite these findings, ABA levels typically do not rise during the natural onset or progression of senescence in temperate deciduous trees (Kane and McAdam, 2023; Zhang et al., 2020). The link between ABA and accelerated senescence may explain why under drought leaves senesce earlier (Frei et al., 2022; Ma et al., 2024; Salleo et al., 2002; Vander Mijnsbrugge et al., 2025). Drought-deciduous species offer further support for this idea, as leaf senescence in these species is closely associated with declining leaf water potential (Ψ_L_) (Brodribb and Holbrook, 2003).

Temperate deciduous trees are especially well-known for their annual leaf shedding each autumn a phenomenon even noted by the Greek philosopher Theophrastus in his third book *Enquiry into Plants*, where he distinguished between wild deciduous and evergreen species (Theophrastus, c.287 BCE). The process of abscission typically begins at the point of organ attachment such as between the petiole and stem, called the abscission zone (AZ), weeks or even months before separation (André et al., 1999). The AZ generally consists of small, densely packed cells arranged in layers, and forms through secondary cell division prior to separation (Sexton and Roberts, 1982), although there is considerable diversity in the nature of AZ development. For instance, *Impatiens* leaves can abscise without forming an AZ layer, and plants such as poinsettia, cotton, and pepper can be induced to abscise before a compressed AZ develops (Gawadi and Avery, 1950). Leading Gawadi and Avery (1950) to speculate that the AZ may serve primarily as a protective barrier post-abscission, rather than simply as a structural weak point for detachment, though this theory has not gained widespread acceptance. Activation of the AZ follows formation during which cells near the attachment site begin producing cell wall-degrading enzymes such as cellulases and pectinases, which weakens tissue cohesion (Abeles, 1969; Arteca, 1996; Bonghi et al., 1992; Mishra et al., 2008; Morre, 1968). This degradation may be accompanied by programmed cell death, especially on the distal (organ-side) portion of the AZ, which further reduces the structural integrity of the attachment site (Pautot et al., 2025). Often occurring simultaneously with AZ activation is the formation of a suberized and lignified protective layer (Arteca, 1996; Gawadi and Avery, 1950; Pautot et al., 2025) at the proximal point of attachment facilitating shedding and sealing of AZ (Arteca, 1996; Crosse, 1951; Hewitt, 1938; van Nocker, 2009). Once the AZ becomes competent to abscise, the final separation occurs. Separation involves the complete physical detachment of the organ, requiring the severing of the remaining tissue, including the xylem, which is often the last physical connection point (Addicott, 1982; Webster, 1968). Separation can be active, where secondary growth at the separation layer physically pushes the organ away (Sexton and Redshaw, 1980), or through turgor-driven ejection, where proximal cells absorb water and lyse forcibly ejecting the organ (Galstyan and Hay, 2018; Osborne, 1989; Sexton, 2001). The exact mechanism of active separation varies by species and organ type (Addicott, 1982). The final separation can also occur via abiotic factors, such as wind, snow, or the weight of the organ itself (Addicott, 1982; Sexton and Redshaw, 1980). Some observations suggest that abiotic factors alone are not always sufficient, and that additional biotic factors, internal or external may be necessary to complete separation (Sexton and Redshaw, 1980; von Wiesner, 1871)

Marcescence refers to the retention of dead leaves on a plant through autumn and winter commonly observed in species of *Quercus* (oaks), *Carpinus* (hornbeams), and *Fagus* (beeches), amongst others (Berkley, 1931). Unlike typical deciduous behavior, in which leaves are shed, marcescent leaves remain attached until spring. An anatomical study of *Quercus palustris* and *Q. coccinea* found that although an AZ developed in both species, it did not activate until April, at that time, cells in the AZ exhibited signs of dissolution (Hoshaw and Guard, 1949). During winter many marcescent leaves are not actively shed but are instead mechanically broken off by wind or snow (Hoshaw and Guard, 1949). This poses a possible risk for wind damage in the winter due to higher surface area (Karban and Pearse, 2021). The ecological significance of marcescence remains poorly understood, one theory suggests that marcescent leaves may aid in nutrient management by creating habitats for animals that fertilize the soil beneath the tree (Kalm, 1770). Another idea is that retaining leaves until spring, when microbial activity and decomposition are higher, allows for more effective recycling of immobile nutrients near the tree (Otto and Nilsson, 1981). Marcescence has also been proposed as a deterrent to winter browsing by deer, making the plant less palatable or accessible (Svendsen, 2001). An additional hypothesis is that extended leaf retention may prolong autumn photosynthesis or provide more time for nutrient resorption, making senescence more efficient (Abadía et al., 1996). In the order Fagales, it has been suggested that marcescence may represent an incomplete evolutionary transition from evergreen to fully deciduous leaf habits (Heberling and Muzika, 2023; Karban and Pearse, 2021).

While there is considerable diversity in the nature of leaf AZ formation and competency across angiosperms the mechanistic regulators of AZ formation have largely focused on floral and fruit abscission, particularly in herbaceous model species such as Arabidopsis and tomato. Ethylene is the most well-described hormone involved in abscission since observations of gas lamp emissions caused premature leaf drop in urban trees (Bakshi et al., 2015; Neljubow, 1901). Ethylene promotes the formation and activation of AZs, by inducing the production of cell wall-degrading enzymes (Agustí et al., 2008; Botton and Ruperti, 2019; Brown, 1997; Mishra et al., 2008; Zhang et al., 2025). Ethylene release is the primary mode of action for agricultural defoliants like Thidiazuron (Suttle, 1985; Zhang et al., 2025). In contrast, auxin is considered a key inhibitor of abscission and an antagonist of ethylene action. A widely accepted theory suggests that as long as there is a high auxin gradient, with more auxin in the distal (leaf) portion than in the proximal (stem) region, abscission is suppressed (Addicott et al., 1955; Dong et al., 2021; Hall, 1952; Liu et al., 2022; Louie and Addicott, 1970; Osborne, 1989). Although studies in deciduous trees are limited, research on hybrid poplars supports the role of auxin as an abscission inhibitor (Jin et al., 2015). Ethylene exposure is thought to reduce auxin transport (Riov and Goren, 1979), while lower auxin levels in the AZ increase sensitivity to ethylene (Taylor and Whitelaw, 2001). Although changes in auxin levels can trigger AZ activation and cellular degradation, in ethylene-insensitive Arabidopsis flowers (Bleecker and Patterson, 1997; Jin et al., 2015). Other hormones play more minor roles in AZ formation. Cytokinins may reduce tissue sensitivity to ethylene and slow leaf senescence (Gomez, 2011; Xu et al., 2019). While ABA, originally named for its proposed role in abscission in cotton (Davis and Addicott, 1972; Ohkuma et al., 1965, 1963), may only contribute to drought-induced abscission by promoting stomatal closure, which reduces photosynthesis (Downton et al., 1988) and auxin production (Avery Jr et al., 1937; Osborne, 1973). ABA can also stimulate ethylene production (Cracker and Abeles, 1969) this may be a feedback from ABA promoting early senescence (Arteca, 1996; Kane and McAdam, 2023) and chlorophyll degradation which can lead to ethylene production (Ito et al., 2022).

Beyond an active-metabolic regulation of abscission, a role of xylem embolism in triggering leaf shedding has been proposed. Under water stress, tension builds within the xylem water column, and if drought becomes severe enough, xylem embolism can form leading to tissue damage or even whole-plant death (Kaack et al., 2021; Tyree and Sperry, 1989). To mitigate such risks, some plants exhibit hydraulic segmentation, where downstream tissues (leaves) are more vulnerable to drought than upstream tissues (stems and roots) (Cardoso et al., 2022; Charrier et al., 2016; Guan et al., 2022). This anatomical and physiological feature underpins the hydraulic fuse hypothesis, which proposes that leaves act as “hydraulic fuses”, the first tissues to fail during drought, thereby protecting stems by shedding leaves in response to embolism formation (Tyree et al., 1993; Wolfe et al., 2016). Support for this hypothesis comes from studies showing that in both drought- and winter-deciduous species, leaf senescence and abscission often coincide with reductions in leaf and/or stem hydraulic conductance. These declines are typically associated with embolism and, in some cases, xylem blockage by tyloses, structures that occlude xylem vessels (Brodribb et al., 2002; Machado and Tyree, 1994; Pallardy and Rhoads, 1997; Salleo et al., 2002; Sobrado, 1986; Tyree et al., 1993). Salleo et al. (2002) found that both stem and leaf hydraulic conductance declined prior to leaf shedding in *Castanea sativa*, primarily due to embolism and tylosis formation. They proposed that significant stem embolism occurring in late summer may initiate autumn leaf senescence and winter dormancy. Other studies have shown that embolism often forms in leaves before stems, with leaf abscission following embolism development (Brodribb et al., 2002; Hochberg et al., 2017). Moreover, leaf shedding in response to severe drought has been observed across a wide range of plant species, regardless of typical leaf habit, whether evergreen, winter-deciduous, or drought-deciduous (Brodribb and Holbrook, 2005; Dallstream and Piper, 2021; Machado and Tyree, 1994). Despite growing evidence, many questions remain about the causal role of embolism in senescence and abscission.

Due to the limited data available on abscission zone competency (AZC) in woody species and the destructive anatomical nature of most techniques to quantify AZ development and competency we sought to develop a method that could quantify the number of leaves that are competent to abscise before the final physical separation has occurred. Using this method we aimed to better understand the relationship between leaf senescence and abscission, with a focus on how these processes are both overlapping and independently regulated resulting in different deciduous mechanistic types. We measured leaf gas exchange, chlorophyll content, Ψ_L_, foliar ABA concentration, and AZC in four temperate tree and shrub species exhibiting diverse leaf habits during the autumn of 2020. These included two deciduous species, *Ginkgo biloba* L. (Ginkgoaceae) and *Phellodendron amurense* Rupr. (Rutaceae); one brevi-deciduous species, *Lonicera x purpusii* Rehder. (Caprifoliaceae), which is leafless for only a few weeks in late winter (Brailko and Gubanova, 2014; Kane and McAdam, 2024); and one marcescent species, *Quercus falcata* Michx. (Fagaceae), which retains most of its leaves through the winter. We hypothesized that the two fully deciduous species would exhibit similar physiological patterns, including nearly complete chlorophyll degradation and cessation of gas exchange prior to the activation of their AZs. We expected the brevi-deciduous species to follow a similar trajectory, but with senescence-related declines either occurring later or progressing more slowly. For the marcescent species, we predicted that leaves would be killed by frost while retaining a high proportion of their chlorophyll content. In addition to these physiological observations, we also present a novel and accessible method for estimating AZ competency in deciduous species an approach that may improve our understanding of the timing and regulation of leaf drop across diverse plant strategies.

## Materials and methods

### Plant material

All trees and shrubs were established individuals growing on the campus of Purdue University, West Lafayette, IN, USA near Lilly Hall of Life Sciences (N 40.233208, W 869167627. All plants received 15:3:3 N:P:K fertilizer twice a year with no additional irrigation and no pruning for at least three years before measurements were made. Measurements began in early September 2020 and continued every 2-3 days until all leaves had fully abscised (*P. amurense* and *G. biloba*) or were dead on the tree (*Q. falcata* and *L. x purpusii*).

### Measurements

All collections were conducted between 11:00h and 13:00h on full sun or mostly sunny days. To measure CO_2_ assimilation and stomatal conductance an inferred gas analyzer (LI-6800 Portable Photosynthesis System; LI-COR Biosciences, Lincoln, NE, USA) was used. Ambient air from a buffer drum was used for the gas analyzer. Conditions in the cuvette were set to ambient vapor pressure difference between the leaf and atmosphere (VPD). Light conditions were set to a saturating 1500 µmol quanta m^−2^ s^−1^. As soon as assimilation and stomatal conductance was recorded the leaf was wrapped in a damp paper towel and detached from the tree and double bagged in Ziploc sandwich bags (SC Johnson, MI, USA) to minimize water loss during transportation. Leaves were then placed in a dark bag for a minimum of 5 minutes to allow Ψ_L_ to equilibrate. Leaves were then taken back to the lab where Ψ_L_ was measured using a Scholander pressure chamber (PMS Instrument Company, OR, USA). After quantification of Ψ_L_ leaf tissue was harvested into separate tubes for quantification of foliage ABA and chlorophyll content.

Foliage ABA was measured using physicochemical methods with an added internal standard. A subsample of tissue from the middle of the lamina, avoiding the midrib, was taken from each leaf, or the terminal pinna of the imparipinnate leaves of *P. amurense*, for hormone and pigment analysis. The mass of the fresh leaf sample was recorded (OHAUS Corporation, Parsippany, NJ, USA), and then the tissue was covered in –20 °C 80% methanol in water (v/v) containing 250 mg l^−1^ butylated hydroxytoluene (BHT), chopped into fine pieces, and stored in a –20 °C freezer overnight. Extraction in methanol allows for extraction of both free and bound ABA (Georgopoulou and Milborrow, 2012). Leaf tissues were thoroughly homogenized after which 15 ng of deuterium-labeled [^2^H_6_]-ABA (OlChemIm, Olmouc, Czech Republic) was added to each sample before extracting overnight at 4 °C. A 5 ml aliquot of supernatant was taken from each sample and completely dried in a vacuum sample concentrator (Labconco, MO, USA). The sample was then resuspended in 200 μl of 2% acetic acid in water (v/v). The new suspension was then centrifuged at 14 800 rpm for 4 min, and a 100 μl subaliquot was taken for quantification of ABA and internal standard levels using an Agilent 6460 series triple quadrupole LC/MS (Agilent, CA, USA) (McAdam, 2015). After quantification, sample tissue was dried at 70°C in the original tube. Approximate leaf dry mass was quantified by subtracting the mass of the clean empty tube from the mass of the tube containing the dried homogenized leaf material.

Fresh mass was recorded for the second leaf sample harvested similarly for chlorophyll quantification. Leaf tissue was covered with –20 °C acetone containing 250 mg l^−1^ BHT and roughly chopped and stored overnight at –20 °C before being homogenized and extracted overnight at 4 °C. A 100 μl aliquot of supernatant was taken for Chl *a* quantification using an Agilent 1100 high-performance liquid chromatograph, with an Agilent 1100 G1315B diode array detector for UV-visible spectrophotometric detection of pigments (Agilent, CA, USA). A Zorbax StableBond S8-C18 column (4.6 mm×150 mm with 5 µm particle size) (Agilent) and a quaternary pump (Agilent) was used according to McAdam et al. (2022). Chlorophyll *a* (Chl *a*) content was calculated in the injected sample using a standard curve of known quantities of Chl *a.* Chl *a* for the whole sample was then calculated by multiplying this number by the total volume of acetone used to extract the pigment. Total volume of acetone was measured gravimetrically by obtaining the mass of the total homogenized sample in the acetone, then drying the sample at 70°C and re-massing the sample when completely dry. The volume of acetone was calculated by multiplying the mass change and the density of acetone at standard atmospheric pressure at 180m above sea level. Dry mass of the samples was determined by cleaning and redrying the tube and re-massing the clean tube. The mass of the clean empty tube was subtracted from the mass of the tube containing dry matter.

### Measuring abscission zone competency

AZC was determined as the percentage of total leaves that had fallen after rapid freezing and thawing. To assess AZC each time other measurements were taken a neighboring branch was collected and gently placed in a 50 l black plastic bag containing damp paper to minimize water loss. Collected branches typically had between 10-40 leaves. Branches were then placed in a −10°C walk in freezer and left for 2 hours to ensure they had completely frozen. After 2 hours the branches were removed from the freezer and allowed to reach room temperature for 30 minutes. After thawing each branch was gently moved in an arc for 30 seconds to allow any leaves with competent abscission zones fall from the stem. The number of leaves fallen (including those that fell off in the bag while in the freezer) and still attached were counted.

### Vulnerability curves

In *Q. flacata*, and *L. x purpusii* leaf vulnerability curves were measured using the optical method (Brodribb et al., 2016; Cardoso et al., 2022). Vessel length was measured via air injection to determine the longest vessel and avoid open vessel artifacts. Branches at least twice as long as the longest vessel were cut and recut underwater for transport to the lab. Leaves were mounted to dissecting microscopes with transmitted light and imaged every 5 minutes until the leaves were fully desiccated. Water potential was measured using PSY1 stem psychrometer (ICT International) every 10 minutes and checked against Ψ_L_ using a Scholander pressure chamber at 5 points during the dry down. Images were processed using via image subtraction on FIJI (NIH) according to opensourceov.org. Previously published optical vulnerability curves for the same individual of *P. amurense* was taken from Avila et al. 2021.

Leaves of *G. biloba* were too thick to allow the transmitted light needed to visualize embolism using the OV method, instead leaf hydraulic measurements (K_leaf_) were made according to Brodribb and Cochard (2009). Shoots from the measured plant were cut and bench-dried, and K_leaf_ was determined at intervals of 0.1 MPa (Ψ_L_) by measuring the rehydration flux of water into leaves, data was logged on a CR850: Measurement and Control and Datalogger (Campbell Scientific, UT, USA). P_50_ was determined by fitting the relationship between Ψ_L_ and K_leaf_ with a 3 parameter-sigmoidal relationship from the pooled data (n=65 leaves).

### Data analysis

For *P. amurense*, *G. biloba*, and *L. x purpusii* AZC through the season was fitted with a two-segment linear regression to find the break point and significance was tested via a Davies test according to Manandhar et al. (2024). The AZC data for *Q. flacata* was fitted with a single liner regression AZC=-9.8484+DAS*0.1274.

General additive models (GAMs) with standard errors (SEs) were fitted relationships between AZC and other data including CO_2_ assimilation rate, Chl *a* content, foliage ABA content, and Ψ_L_ a GAM with p<0.05 was considered significant.

## Results

### Rate of achieving abscission zone competency varies across leaf habit

In three of the four species measured we saw considerable development of AZC as autumn progressed (Fig. 1). The leaves of the two deciduous species increased in AZC with *P. amurense* and *G. biloba* both achieving complete AZC during our measurements around 134 ± 1.5 days after the summer solstice (DAS) and 133 ± 1 DAS (Fig. 1), respectively. The brevi-deciduous *L. x purpusii* only achieved 67.92 ± 2.7% AZC before having the remaining living foliage killed by severe frost 248 DAS, over 110 days after the two deciduous species had reached maximum AZC (Fig. 1C). In contrast the marcescent *Q. falcata* never had more than 9.13 ± 2.95% AZC by 149 DAS with many individual branches have no AZC when the canopy was killed by a frost (Fig 1D).

**Figure 1.**
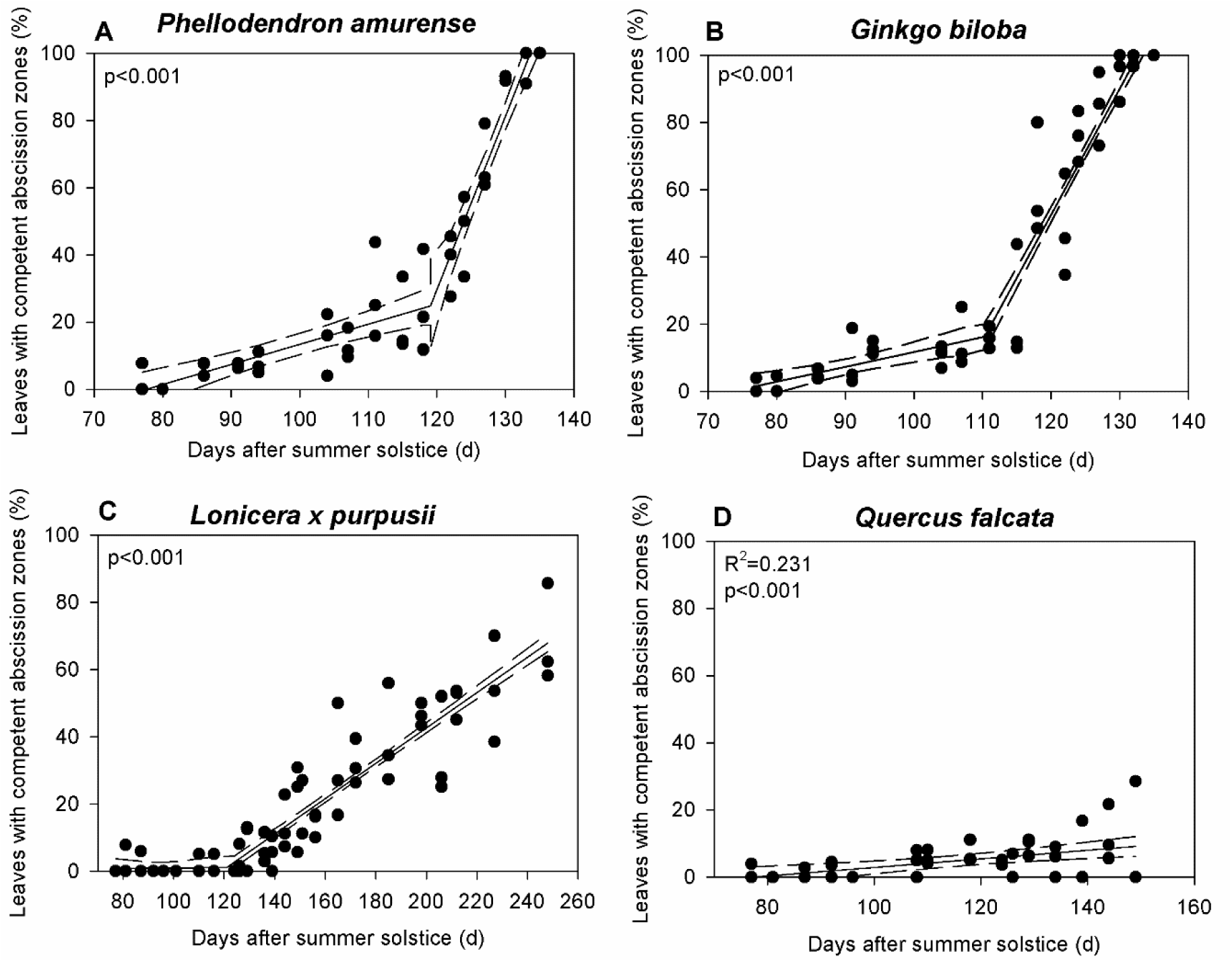
Percentage of leaves with competent abscission zones from early-September 2020 until total defoliation or leaf death when still attached to the tree in the deciduous species *Phellodendron amurense* (A) and *Ginkgo biloba* (B), brevi-deciduous *Lonicera x purpusii* (C). Solid lines depict a segmented regression with p-values according to a Davies’ test shown, dashed lines demarcate the SE of the model. Marcescent *Quercus falcata* (D) the solid line indicates the linear regression with dashed lines showing SE with R^2^ and p-value. Each point represents data from an independent branch containing ten or more leaves.

The two deciduous species appeared to go through two distinct phases where AZC developed slower early in the autumn and then much more rapidly towards the end of the season. *G. biloba* went from 1.57 ± 3.77% AZC on 77 DAS to 16.2 ± 6.4% AZC 33 days later with the rate of AZC increasing by around 4.4‰ per day during this period, however after the break point (110.3±4.27 DAS) the rate increased 7-fold to around 37.6‰ per day with branches approaching complete AZC by 133 DAS (Fig. 1B). Similar trends were observed in *P. amurense* in which AZC in branches increased from 0 ± 5.13% on 77 DAS to 24.83 ± 4.5% on 119 DAS, with AZC increasing at a rate of 6‰ per day (Fig. 1A). After the breakpoint (119.04 ± 2.3DAS) the rate of AZC increased in *P. amurense* to 51.6‰ per day with plants reaching complete AZC by 134 DAS (Fig. 1A). In the brevi-deciduous *L. x purpusii* there was a very minimal increase in AZC early in autumn with AZC increasing from 8.7 ± 23.1‰ on 77 DAS to 1.15 ± 4.45% by 121 DAS representing an increase of only 0.065‰ per day after this the rate increased but slower than in the other deciduous species increasing to 67.92 ± 2.7% 248 DAS showing an increase of 5.3‰ per day (Fig 1C). We observed a lack of trend in *Q. falcata* with no significant change in AZC between 77 DAS and 149 DAS when leaves were killed by frost (Fig. 1D).

### Diversity in chlorophyll reclamation and productivity decline

We observed a diversity of responses in leaf Chl *a* content as AZC increased during the autumn across the four species sampled (Fig. 2). In the deciduous species we observed a rapid decline in Chl *a* content when AZC was less than 20%. In *P. amurense* Chl *a* content in leaves declined from an average of 93.11 ± 5.75 % of max Chl *a* content when there was no AZC to 57.26 ± 4.06 % when there was 20% AZC when AZC increased after this point Chl *a* content decline slowed dropping by only another 14.97 ± 4.32% to 42.29 ± 6.38% by the time of complete AZC in *P. amurense* (Fig. 2A). *G. biloba* had a similar sharp decline in Chl *a* content early in AZC declining from an average of 90.43 ± 5.33 % when there was 1.28% AZC on branches to 39 ± 5.63% when AZC had increased to 20% (Fig. 2B). Chl *a* in *G. biloba* declined a further 24.52 ± 7.52% to 14.48 ± 6.14% by the time there was complete AZC (Fig. 2B). In the brevi-deciduous *L. x purpusii* high levels of Chl *a* content were maintained in leaves for most of autumn until leaves were killed by frost in winter (Fig. 2C). In *L. x purpusii* Chl *a* declined from 80 to 65% of maximum levels as AZC increased from 0 to 60% before sharply declining to 43.55 ± 11.78% of maximum as AZC increased to 68.74% at which point the remaining leaves were killed by frost (Fig. 2C). *Q. falcata* experienced a rapid and continuous decline in Chl *a* across only small increases in AZC (Fig. 2D). Declining from 75.76 ± 11.11% of maximum to none as AZC increased to only 12% by the time the leaves were killed by frost (Fig. 2D).

**Figure 2.**
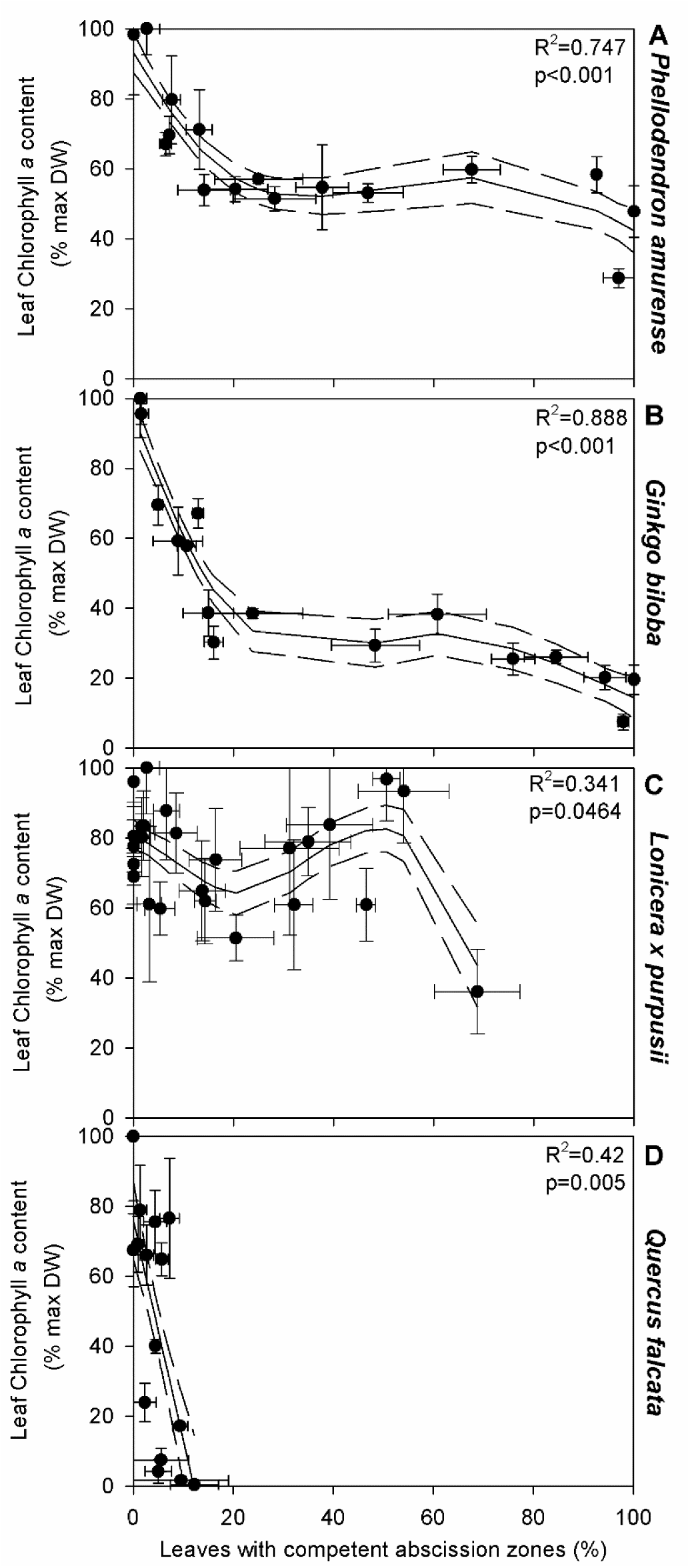
The relationship between leaf chlorophyll a content expressed as a percentage of maximum and the percentage of leaves with competent abscission zones in the deciduous species *Phellodendron amurense* (A) and *Ginkgo biloba* (B), brevi-deciduous *Lonicera x purpusii* (C) and marcescent *Quercus falcata* (D). Solid lines depict a general additive model with R^2^ and p values shown, dashed lines demarcate the SE of the model. Each point represents an average (n=3) of data collected on a single day with bi-directional error bars showing SEs.

In both the deciduous and brevi-deciduous species we observed declines in leaf CO_2_ assimilation rates as AZC increased in the autumn but saw no significant relationship between CO_2_ assimilation and AZC in the marcescent *Q. falcata* (Fig. 3). *P. amurense* and was the only tree to maintain assimilation rates above 1% of maximum throughout the autumn, maintaining 31.5 ± 12.02% percent of maximum assimilation when AZC had reached 96.97% (Fig. 3A). In contrast both *G. biloba* and *L. x purpusii* both had much lower assimilation rates by the time of complete AZC or when leaves died in frost. In *G. biloba* assimilation rates fell from 62.63 ± 7.62 % of mximum to an average of −0.26 ± 12.08% of maximum once AZC appraoched 97.91% (Fig. 3B). *L. x purpusii* saw similar declines dropping from 74.19 ± 5.27% of maximum when no leaves were competent to abscise to 0.47 ± 11.48% of maximum when AZC had increased to 68.74% (Fig. 3C).

**Figure 3.**
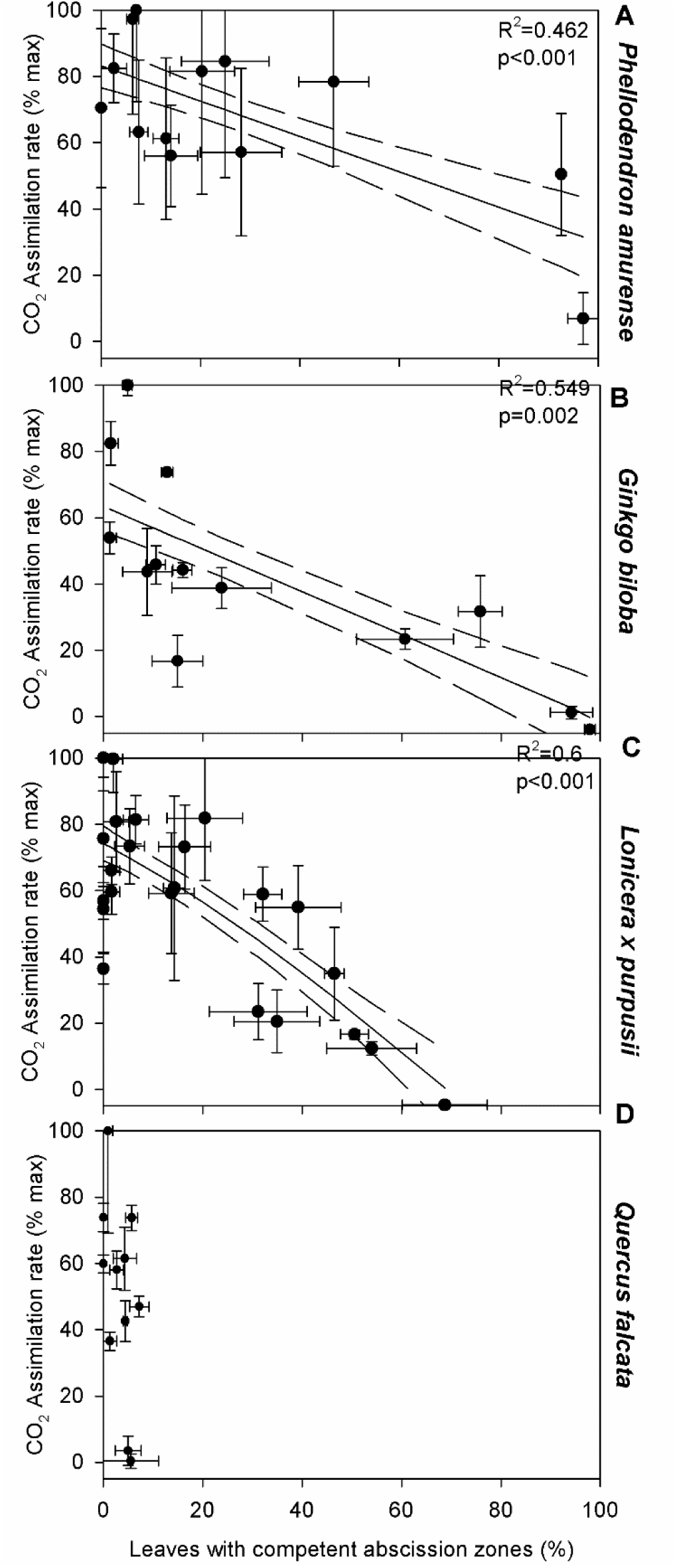
The relationship between CO_2_ assimilation rates expressed as a percentage of maximum and the percentage of leaves with competent abscission zones in the deciduous species *Phellodendron amurense* (A) and *Ginkgo biloba* (B), brevi-deciduous *Lonicera x purpusii* (C) and marcescent *Quercus falcata* (D). Solid lines depict a general additive model with R^2^ and p values shown, dashed lines demarcate the SE of the model. If no line is shown this indicates that the general additive model was not significant. Each point represents an average (n=3) of data collected on a single day with bi-directional error bars showing SEs.

### Leaf water status and ABA during increasing abscission zone competency

In the two deciduous species we observed increasing Ψ_L_ as AZC approached 100%. With *P. amurense* progressing steadily from −1.27 ± 0.04 MPa when there was no AZC to −0.6 ± 0.07 MPA as AZC approached 100% (Fig. 4A). In *P. amurense* foliage ABA level declined from 0.399 ± 0.107 μg g^−1^ DW when there was no AZC to 0.134 ± 0.117 μg g^−1^ DW when AZC had increased to 46% before increasing to 0.97 ± 0.113 μg g^−1^ DW was AZC approached 100% (Fig. 5A). In contrast foliage ABA levels in *G. biloba* rose almost 5 times as AZC increased in the autumn from 0.489 ± 0.2 μg g^−1^ DW to 2.34 ± 0.257 μg g^−1^ DW (Fig. 5B), during this same period *G. biloba* leaves became more hydrated rising from −1.22 ± 0.08 MPa to −0.37 ± 0.1 MPa (Fig 5B). In *L. x purpusii* we saw declining foliage ABA levels from 0.284 ± 0.056 μg g^−1^ DW to 0.08 ± 0.607 μg g^−1^ DW as AZC increased from none to 68% (Fig. 5C), we observed no significant relationship between AZC and Ψ_L_ (Fig. 4C). In the marcescent *Q. falcata* we observed Ψ_L_ remained relatively steady rising slightly from −1.52 ± 0.31 MPa to −0.93 ± 0.41 MPa between no AZC and 9% before declining sharply to −4.22 ± 0.45 MPa as AZC increased 12% and leaves were killed by frost (Fig. 4D). During this time foliage ABA levels rose from 0.125 ± 0.535 μg g^−1^ DW to 3.48 ± 0.86 μg g^−1^ (Fig. 5D). During our measurement period Ψ_L_ did not decline sufficiently to cause leaf embolism or major declines in leaf hydraulic conductance (Fig 5A and B). In *P. amurense* the water potential at P50 was −2.54 ± 0.29, in G. biloba it was −2.46 ± 0.13 (Fig 4A and B). In *L. x purpusii* the lowest recorded mean Ψ_L_ was −2.91 ± 0.6 MPa corresponding to approximately 40% leaf embolism, with a mean leaf P50 in this species of - 3.31 ± 0.17 MPa (Fig. 4C). In *Q. falcata* Ψ_L_ declined to −4.34 ± 1.9 MPa which was sufficient to induce at least 35% embolism in the leaf, with a mean leaf P50 in this species of −5.28 ± 0.4 MPA (Fig. 4D)

**Figure 4.**
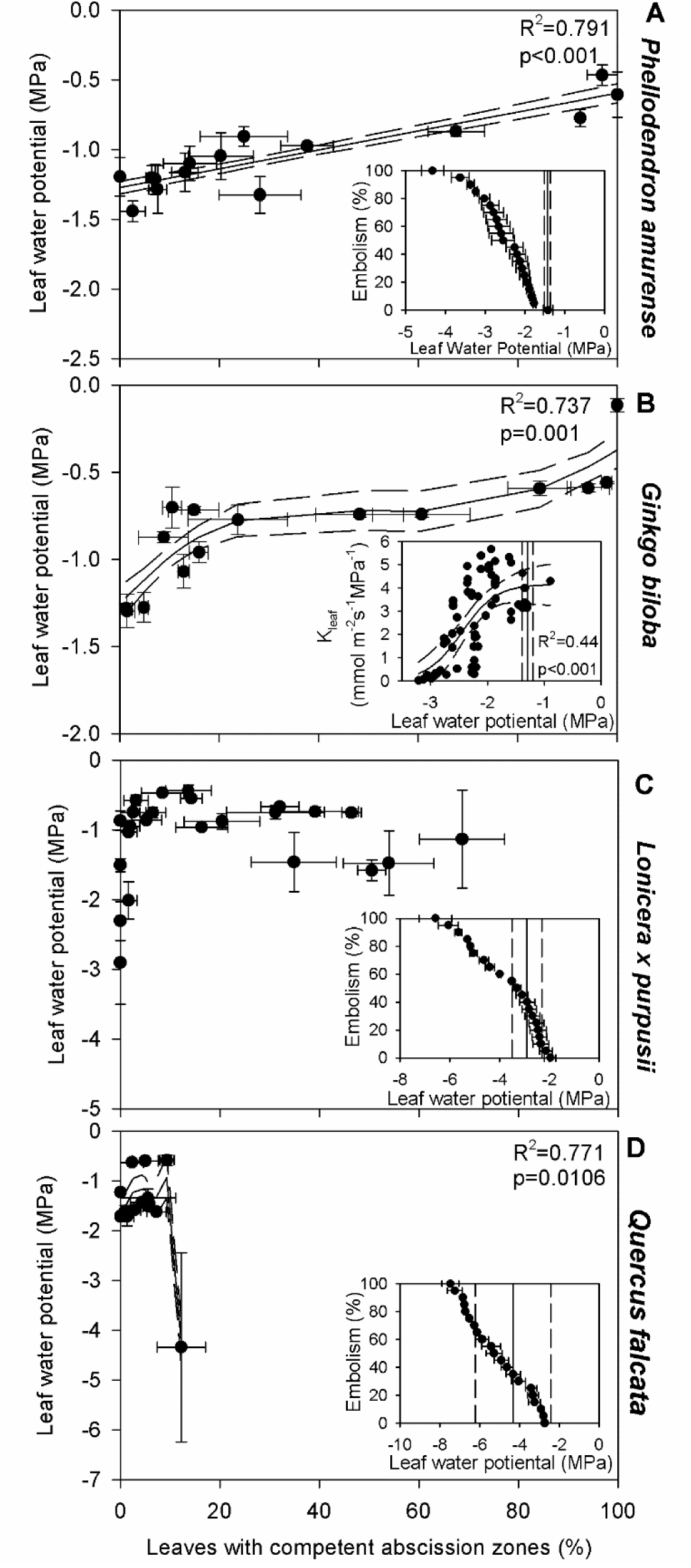
The relationship between leaf water potential and percentage of leaves with competent abscission zones in the deciduous species *Phellodendron amurense* (A) and *Ginkgo biloba* (B), brevi-deciduous *Lonicera x purpusii* (C) and marcescent *Quercus falcata* (D). Solid lines depict a general additive model with R^2^ and p values shown, dashed lines demarcate the SE of the model. If no lines are shown this indicates that the general additive model was not significant. Each point represents an average (n=3) of data collected on a single day with bi-directional error bars showing SEs. Inserts depict leaf average vulnerability (n=3) with SE to embolism using the optical method *Phellodendron amurense* (A), *Lonicera x purpusii* (C), and *Quercus falcata* (D) and the hydraulic method *Ginkgo biloba* (B), the solid line shows a three-parameter sigmoidal fit with SE shown in dashed lines. The solid vertical lines show the most negative leaf water potential observed during the collection period with SE show with vertical dashed lines. Vulnerability data from *Phellodendron amurense* (A) comes from Avila *et al*. (2021).

**Figure 5.**
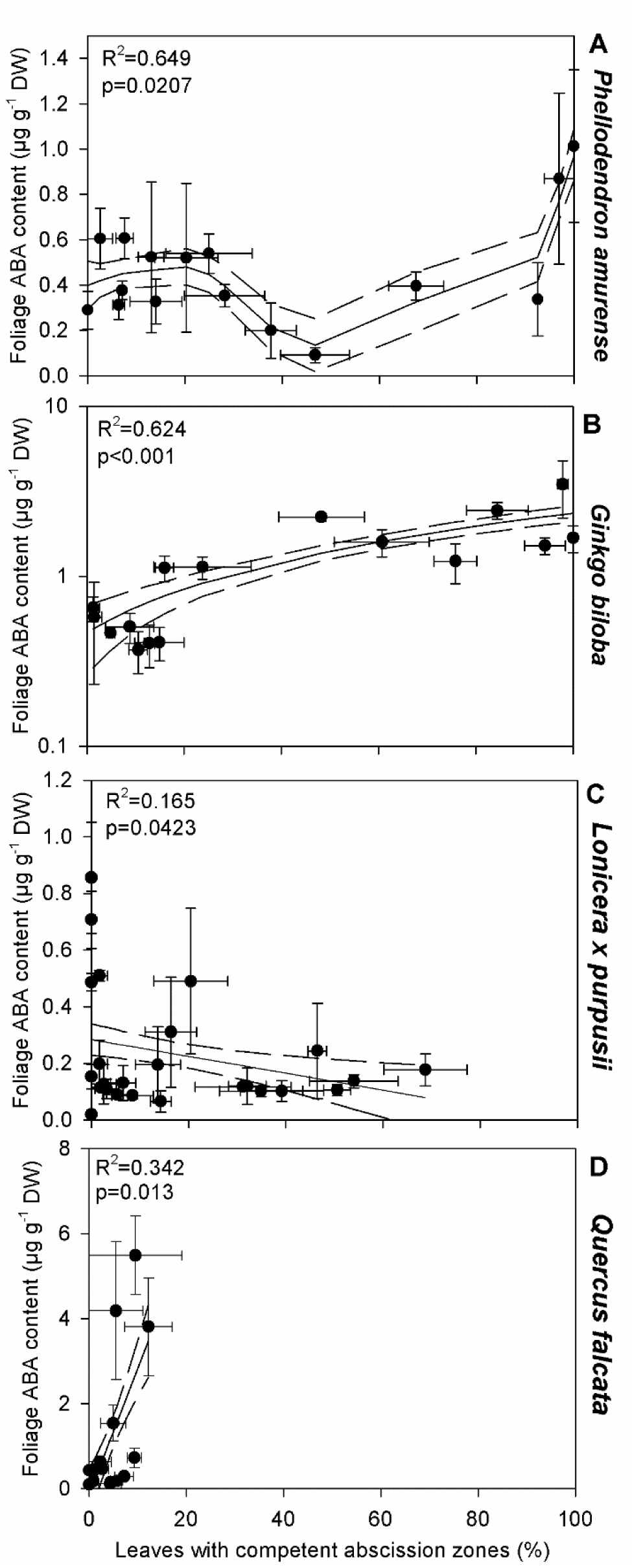
The relationship between foliage ABA content and the percentage of leaves with competent abscission zones in the deciduous species *Phellodendron amurense* (A) and *Ginkgo biloba* (B), brevi-deciduous *Lonicera x purpusii* (C) and marcescent *Quercus falcata* (D). Solid lines depict a general additive model with R^2^ and p values shown, dashed lines demarcate the SE of the model. Each point represents an average (n=3) of data collected on a single day with bi-directional error bars showing SEs.

## Discussion

### Abscission zone competency is a major driver of temperate leaf functional types

Our results show that a direct driver of leaf habit in temperate species is the rate at which abscission zones become competent. Using a novel freezing method, we demonstrate that AZC develops at different rates across deciduous species, and more gradually in brevi-deciduous species (Fig. 1). In the marcescent *Q. falcata*, which retained most of its dead leaves, we observed very little AZC throughout the study period (Fig. 1D). This aligns with previous anatomical observations in marcescent oaks (Hoshaw and Guard, 1949). The freezing method confirmed that leaves without AZC did not separate, validating that our technique does not artificially induce abscission. Therefore, *Q. falcata* served as an effective control in establishing the utility of our method for quantifying AZC. We believe this simple and accessible freezing technique has potential to advance understanding of AZC development in deciduous trees. Further testing is required to assess its applicability of this method across other types of deciduousness, such as in drought-deciduous species and herbaceous plants.

In the deciduous species *P. amurense* and *G. biloba*, we observed rapid AZC development during autumn 2020 (Fig. 1). The progression occurred in two phases: an initial slow phase, where AZC increased by only fractions of a percent per day, followed by a later phase with a sharp rise of 3–5% AZC per day. At the onset of this accelerated phase, *G. biloba* had reached ∼16% AZC, while *P. amurense* was at ∼26% (Fig. 1). Interestingly, although *G. biloba* began this second phase 10 days earlier than *P. amurense*, it only reached full (100%) AZC one day sooner. *G. biloba* is widely known for its highly synchronized leaf drop, with most leaves falling within a single day (Hacskaylo and Moke, 1964). This phenomenon has received considerable public attention (Casanave, 2005; Hoplamazian, 2023; Kurzius, 2019; Meyer, 2017), and even inspired a poem by U.S. Poet Laureate Howard Nemerov in 1975, titled *The Consent* (Nemerov, 1981). Decades of annual contests are held for predicting the date *G. biloba* trees will defoliate (Casanave, 2005; Hoplamazian, 2023). These observations have led to speculation that AZC in *G. biloba* is more tightly synchronized than in other deciduous species. However, empirical support for this claim remains limited (Casanave, 2005; Hoplamazian, 2023; Kurzius, 2019; Meyer, 2017). *G. biloba* appears to form and activate AZs in a very similar process and speed to other species including the deciduous *Fraxinus americana* (Facey, 1950; Mosher and Smith, 1970). Both measured deciduous species in our study exhibited similar AZC progression patterns (Fig. 1). We hypothesize that *G. biloba* adopts a conservative strategy initiating AZC earlier than other co-occurring species allowing it to achieve 100% AZC before major frost or wind events. This may explain the uniform leaf drop observed when such events occur, while other species may only partially defoliate. We also propose that final leaf separation in both *G. biloba* and *P. amurense* requires an external physical force, often a hard freeze (<-5°C), in colder climates. This freezing may damage the AZ tissue, such that thawing completes the separation process. More research is needed to explore the diversity in AZC rates across deciduous plants and to understand how many other species similarly develop complete AZ formation before any leaves are shed like in *G. biloba*. It is not known what the mechanism is that causes AZ damage and separation following freezing, but it likely requires severing of xylem tissue.

We have previously shown that *L. x purpusii* can successfully maintain photosynthesizing leaves into late winter (February in North America) (Kane and McAdam, 2023, 2024). We speculate that this capacity for cold tolerance maintaining function down to −20°C is a key factor in the brevi-deciduous leaf habit (Kane and McAdam, 2024). Other *Lonicera* spp. naturalized in Eastern North America also exhibit this trait, surviving mild frosts in both spring and autumn. This allows them to extend growing seasons, outcompeting native herbs and shrubs by accumulating more carbon during periods when canopy trees are leafless (Fridley, 2012; McEwan et al., 2009; Smith, 2013). This prolonged growing period is likely supported by delayed AZC. *L. x purpusii* showed significantly slower AZC development compared to the strictly deciduous *P. amurense* and *G. biloba*. Whether this delay results from reduced sensitivity to environmental cues such as day length and temperature, or from physiological feedback related to continued photosynthesis at low temperatures, remains an open question.

### Leaf chlorophyll and productivity during leaf abscission

Our results from *P. amurense* and *G. biloba* reveal similar patterns in the relationship between Chl *a* content and AZC. In both species, Chl *a* declined rapidly until AZC reached approximately 20%. Beyond this point, Chl *a* levels plateaued, resulting in 42% and 14% of chlorophyll remaining in the abscised leaves of *P. amurense* and *G. biloba*, respectively (Fig. 2). These findings suggest that both species undergo two partially overlapping phases: an initial phase of senescence, during which chlorophyll is actively resorbed, followed by a transition in which senescence halts as AZC increases rapidly past 20%. Although both species followed this trend, *G. biloba* was smore efficient at degrading chlorophyll, indicating a potentially greater reabsorption of nutrient reserves from leaves (Fig. 2) (Bhat et al., 2019). This greater chlorophyll degradation may also influence leaf litter quality and decomposition rates (Bhat et al., 2019; Sanaullah et al., 2010). The changes in chlorophyll content are mirrored by the patterns we observed in CO₂ assimilation. While *P. amurense* continued photosynthesizing at 31% of its peak rate even when over 95% of its leaves had become competent to abscise, *G. biloba* had completely ceased CO₂ assimilation by that same point (Fig. 3). This phenomenon where senescence halts or remains incomplete while abscission progresses is not unprecedented. For instance, under stress conditions like dehydration, leaves may be shed with most chlorophyll still intact (Osborne, 1973). Similarly, in stay-green soybean mutants, senescence is significantly delayed or halted while abscission proceeds normally (Guiamét et al., 1990). Fuente and Leopold (1968) proposed that abscission is not necessarily a consequence of senescence but rather that senescence can accelerate abscission by increasing the sensitivity of the AZ to ethylene leading to faster development of AZC. Phloem function is critical in this context, as it serves as the primary route for nutrient resorption from senescing leaves (Eschrich et al., 1988). If the formation or activation of the AZ physically disrupts phloem tissue, nutrient transport may be impaired. Increased callose deposition possibly part of the formation of the protective layer in the AZ (Jaffe and Goren, 1988), could further inhibit nutrient remobilization if AZ development outpaces leaf senescence. More anatomical and physiological work is needed to understand the roles of phloem disruption and callose deposition in limiting full nutrient resorption from leaves during abscission.

In *L. x purpusii,* high Chl *a* levels were maintained until AZC reached 54%. After this point, Chl *a* content declined from 80% to 43% as AZC progressed to 68% (Fig. 2). Despite retaining high chlorophyll levels, CO₂ assimilation decreased steadily as AZC increased. This suggests that chlorophyll was preserved even as photosynthetic activity waned, likely due to decreasing temperatures as winter approached. The ability to maintain chlorophyll during cold conditions, combined with considerable freezing tolerance (Kane and McAdam, 2024), may allow *L. x purpusii* to continue low levels of photosynthesis on warmer winter days similar to evergreen species (Hughes and Smith, 2007) (Fig. 2 and 3).

An intriguing insight from our data relates to the potential benefits of marcescence. *Q. falcata* was the only species in which Chl *a* dropped to undetectable levels, indicating that it may have been more successful in extracting mobile nutrients like magnesium prior to leaf death. This extended period for nutrient remobilization, supports a key hypotheses about the ecological advantages of marcescence (Abadía et al., 1996) (Fig. 2 and 3).

### Water relations and ABA dynamics

It has previously been hypothesized that xylem embolism or blockage by tyloses may be necessary for leaf abscission due to declines in leaf hydraulic conductance often observed during senescence and prior to shedding (Brodribb et al., 2002; Machado and Tyree, 1994; Pallardy and Rhoads, 1997; Salleo et al., 2002; Scott et al., 1967; Sobrado, 1986; Tyree et al., 1993). Our findings contrast with this theory. Although we did not measure leaf hydraulic conductance or tylose formation, we observed that leaves of both deciduous species became more hydrated as AZC increased, with Ψ_L_ rising by approximately 1 MPa as AZC progressed from 0% to 100% (Fig. 4). This trend is counterintuitive given the concurrent increase in foliar ABA levels with AZC (Fig. 5), since elevated ABA is typically associated with drought conditions, reduced water potential, and turgor loss (Beardsell and Cohen, 1975; McAdam and Brodribb, 2016; Wright, 1977). In *P. amurense*, ABA levels initially declined with increasing AZC but began to rise again between 46% and 100% AZC. In contrast, *G. biloba* showed a continuous increase in ABA as AZC rose (Fig. 5). For *L. x purpusii*, no significant relationship was observed between Ψ_L_ and AZC (Fig. 4), but ABA levels declined as AZC increased (Fig. 5), unlike in the deciduous species. This suggests that increased ABA is not essential for leaf abscission, although it may enhance the process. Supporting this, we previously demonstrated that girdled *L. × purpusii* branches where ABA levels are elevated defoliated nearly 90 days earlier than intact branches (Kane and McAdam, 2023). We also found that while the deciduous species never reached a Ψ_L_ that would have caused embolism to form in the xylem or a decline in K_leaf_, the brevi-deciduous and marcescent species likely reach a Ψ_L_ that could have caused close to or more than 40% embolism (Fig. 4 and 5). Our data contrasts with the idea that embolism is key to autumnal leaf abscission as the only plants that likely embolized either retained dead leaves on the tree (*Q. falcata*) or were retained alive on the plant for another 171 days (*L. × purpusii*) 113 days longer than either deciduous species (Fig. 4)

We propose three possible hypotheses to explain our observations of the relationship between AZC and foliage ABA levels:

**1)** ABA my help activate abscission zones. Although no longer considered the primary driver of abscission, ABA was once thought to be a central hormone in regulating this process (Davis and Addicott, 1972; Ohkuma et al., 1965, 1963). The effect of ABA on abscission zones may be indirect as high ABA levels may induce ethylene production (Cracker and Abeles, 1969) or ABA acting directly on stomata to reduce photosynthesis may hasten senescence (Arteca, 1996; Downton et al., 1988; Kane and McAdam, 2023) by lowering auxin levels in leaves (Avery Jr et al., 1937; Osborne, 1973) and promoting ethylene production (Ito et al., 2022). This mechanism may help explain high leaf shedding rates under drought conditions (Brodribb et al., 2002; Brodribb and Holbrook, 2005; Dallstream and Piper, 2021; Hochberg et al., 2017; Machado and Tyree, 1994; Salleo et al., 2002) when ABA levels typically rise (Daszkowska-Golec, 2016; Kane and McAdam, 2023; Wright, 1977). Support for this hypothesis also comes from a spontaneous ABA deficient mutant of *Betula pubescens* (form *hibernifolia*), which shows a marcescent habit, retaining dead leaves for years, and having lower ABA levels than wild-type trees (Rinne et al., 1992).
**2)** ABA accumulation may result from AZ activation and phloem disruption. Another possibility is that the increase in ABA is caused by physical changes associated with AZ activation, particularly the disruption or blockage of phloem tissues (Jaffe and Goren, 1988), which normally serve as a pathway for exporting foliar ABA and its conjugates (Castro et al., 2019; Jeschke et al., 1997). This disruption is similar to girdling, which is known to raise ABA levels and decouple ABA content from Ψ_L_ (Castro et al., 2019; Dann et al., 1984; Kane and McAdam, 2023; Lihavainen et al., 2021; López et al., 2015; Mitchell et al., 2017; Setter et al., 1980). It has been suggested that phloem flux significantly declines with reduction in assimilation (Sevanto, 2014), but work in other plants has found that phloem loading can continue even after turgor loss point when stomata are completely shut and there is no assimilation (Gersony and Holbrook, 2022).
**3)** ABA increase may be a general marker of impending leaf death. Finally, the rise in ABA could simply reflect a physiological signal associated with leaf senescence and death. We have previously shown that ABA levels spike as leaves approach incipient death across all major vascular plant lineages (McAdam et al., 2022). This increase is seen in leaves undergoing drought, freezing stress, and leaf senescence (sunflower) and this increase was observed in ABA deficient mutants and peaking type drought plants which switch off ABA production during longer term droughts (Brodribb et al., 2014; McAdam et al., 2022; Mercado-Reyes et al., 2024). With this in mind it is possible that the increase in ABA levels may just be related to the impeding death of the leaf possible caused by spontaneous oxidation of ABA precursors by reactive oxygen species (ROS) (McAdam et al., 2017; McAdam et al., 2022) which are known to rise during leaf senescence (Khanna-Chopra, 2012).

Further experimentation will be necessary to distinguish between these hypotheses and fully elucidate the role of ABA in leaf abscission. Understanding these dynamics will improve our broader knowledge of how plants coordinate senescence, nutrient remobilization, and leaf shedding. With the possibility that one or more of these hypotheses may prove true across the plant kingdom.

### Conclusions

As climate change continues to affect phenological events such as leaf-out, senescence, and abscission, it becomes increasingly important to understand the physiological and hormonal mechanisms driving these processes. There is currently an active debate about how climate change will influence autumn phenology in temperate deciduous forests. Some studies suggest that warming temperatures will extend the growing season and enhance carbon sequestration in these ecosystems (Calinger and Curtis, 2023; Grossiord et al., 2022; Menzel and Fabian, 1999; Norby, 2021; Norby et al., 2003). Newer modeling studies propose that these forests may already be near their maximum productivity, and that earlier spring leaf-out could lead to correspondingly earlier senescence and abscission (Zani et al., 2021, 2020; Zohner et al., 2023). Understanding the mechanisms underlying senescence and abscission is therefore critical for improving models of leaf lifespan and seasonal carbon dynamics. To help address this, we present a novel method for quantifying AZC that requires minimal equipment, only a freezer, and provides reliable, direct measurements. Our findings show that in deciduous species, AZC increases rapidly in autumn, typically following a period of declining Chl *a* content. This rise in AZC is accompanied by significant reductions in CO₂ assimilation, increased Ψ_L_, and elevated foliar ABA levels. In contrast, brevi-deciduous species have a much slower rate of AZC development, requiring several months to reach levels comparable to those in the deciduous species. These species also exhibited lower ABA levels as AZC progressed. The marcescent species in our study served as an effective control for our new AZC quantification method. It showed little AZC development over the study period but did display full remobilization of leaf chlorophyll, supporting the method’s validity. Despite these insights, many questions remain about the anatomy and physiology of leaf abscission, particularly in long-lived woody species. Much of the existing literature focuses on abscission in herbaceous plants and reproductive structures such as fruits and flowers. Expanding our understanding of leaf abscission in perennial deciduous species will be essential for predicting how forest ecosystems respond to a changing climate.

## Acknowledgments

We thank The Oak Spring Garden Foundation and their staff, especially James Adelman and the 2025 Bellflowers cohort. We acknowledge the use of the facilities of the Bindley Bioscience Center (National Institutes of Health-funded Indiana Clinical and Translational Sciences Institute), particularly the Metabolite Profiling Facility. We would like to thank Missy Holbrook for helpful discussions about senescence, abscission, and drought and Kean Kane for help with field measurements and Anju Manandhar for helping with segmented regressions.

## Author contribution

SM and CK: Conceptualization; CK collected, analyzed the data, and prepared the manuscript with input and supervision from SM. IR collected and analyzed optical vulnerability curves.

## Data availability

The data supporting the findings of this report are available upon request to the corresponding author CK

